# *genomalicious*: serving up a smorgasbord of R functions for performing and teaching population genomic analyses

**DOI:** 10.1101/667337

**Authors:** Joshua A. Thia

## Abstract

Turning SNP data into biologically meaningful results requires considerable computational acrobatics, including importing, exporting, and manipulating data among different analytical packages and programming environments, and finding ways to visualise results for data exploration and presentation. I present GENOMALICIOUS, an R package designed to provide a selection of functions for population genomicists to simply, intuitively, and flexibly, guide SNP data through their analytical pipelines, within and outside R. At the core of the original GENOMALICIOUS workflow is the conversion of genomic variant data into a `data.table` object. This provides a useful way of storing large amounts of data in an intuitive format that can be easily manipulated using methods unique to this object class. Over time, GENOMALICIOUS has grown to cater to a range of analyses in population structure and demography, adaptive evolution, quantitative traits, and phylogenetics. Researchers using pooled allele frequencies, or individually sequenced genotypes, are sure to find functions that accommodate their tastes in GENOMALICIOUS. The simplicity and accessibility of pipelines in GENOMALICIOUS may also serve as a useful tool for teaching basic population genetics and genomics in an R environment. The source code and a series of tutorials for this package are freely available on GitHub

## INTRODUCTION

The abundance of tools available to modern population geneticists has provided means to answer questions about biodiversity, demography, and evolution in a multitude of ways. Population genomic approaches have enriched our investigations of the biological world by providing data that is abundant (in terms of number of loci) and resolute (in terms of the scale of questions we can address). The declining cost of individual sequencing, the cost efficiency of pooling, and relative ease of assembling reduced representation datasets, is making genomics more affordable and accessible. This has greatly facilitated the use of genomic data to study non-model organisms and wild populations in unprecedented ways.

Choosing to apply genomic information to population genetic questions comes with the challenge of understanding basic computer programming. First and foremost, a grasp of the Unix environment is essential for turning sequence reads into genotype calls. Getting beyond this first hurdle can be exceedingly difficult for first-time genomic data handling. Traversing the proverbial “Valley of Assembling, Calling, Filtering, and Agonising Over Parameter Choices” represents a steep learning curve that can make the choice to use genomic approaches daunting. However, the journey does not end there. The next major task involves decisions on how best to analyse the potential thousands of genome-wide markers obtained from assembly–variant calling pipelines.

The choice of analyses and associated programs depends on an investigator’s hypotheses and personal preferences. Most researchers are likely to end up in R, given its widespread use among scientists (particularly biologists) for statistical analysis and data visualisation. However, many population genetics R packages require very distinct data structures or are designed in a way that may constrain flexibility in how a researcher chooses to handle their data. Furthermore, some desired tools might not be available in R, requiring computational acrobatics to import, export and manipulate data to move among the various programs in a researcher’s analysis pipeline.

Beyond these practical points, there is also a need for an investigator to understand genomic data and population genetic principles to effectively apply genomic approaches to their research. This foundational knowledge may be acquired in different settings, including as a student in a university classroom, or on the job, for example, as a wildlife practitioner or as a postdoc required to incorporate genomics into their work. Supervisors may lack the essential skills and/or knowledge required to offer guidance to their post-graduate students or employees. This can make the application of genomics daunting for first timers and those without population genomics training.

Here, I present GENOMALICIOUS (think “delicious”, but “genoma”, *i*.*e*. “**<u>genom</u>**ic **<u>a</u>**nalyses”), an R package designed to help facilitate the performing and teaching of population genomic analyses. I wanted this package to organise data and pipelines in intuitive ways, increase the accessibility of genomic analyses, and to have minimal/reasonable limitations on data structures. In the sections that follow, I briefly highlight and describe five main degustation menus that GENOMALICIOUS provides and how these can be incorporated into analysis pipelines of population genetic and genomic data.

## IMPORT

The typical datatype a population genomicist will be working with are SNPs recorded as variant calls in a VCF file format. Though packages already exist to import VCF files into R, the returned objects do not necessarily have immediately intuitive structures or lack clear, straightforward ways to manipulate them. Furthermore, some analytical functions take VCF files as their direct input, but this reduces the extent to which manipulation and piping of SNP data can occur directly in R.

In GENOMALICIOUS, I provide a method to import VCF files into R as a long-format `data.table` object. This may not be the most efficient way to handle extremely large datasets, but even for thousands of loci, over hundreds of individuals, this is a useful way to store population genomic data. The function `vcf2DT()` takes VCF files, which are in in wide-format, where each sample has its own column and loci are in rows, and converts them into long-format, samples and loci are both in columns (see Figure 1).

**Figure 1.**
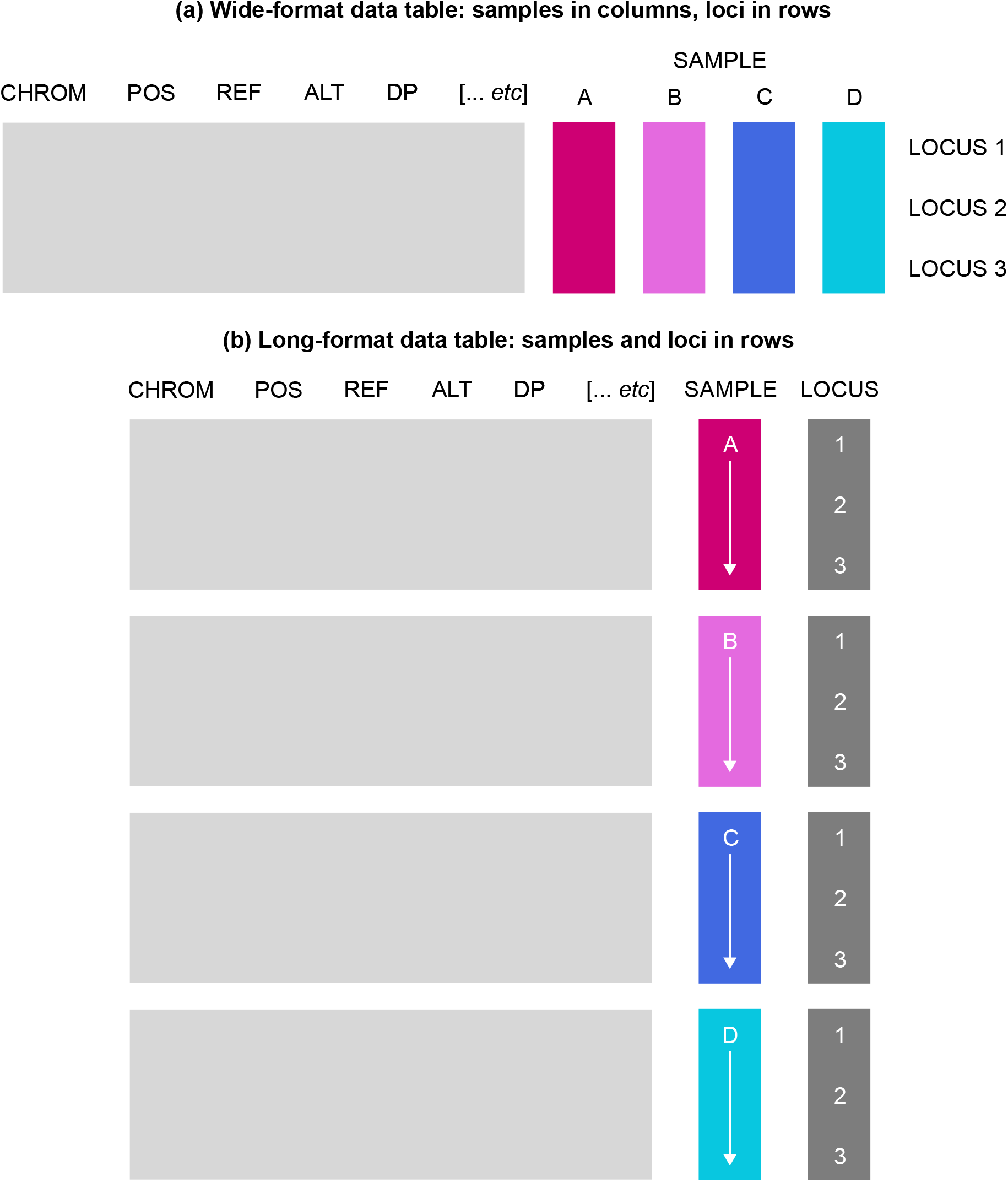
An illustration of wide- and long-format data structures in genetic data. Coloured rectangles represent different data. Metadata is represented by the light grey boxes. Samples are represented by the coloured boxes. Loci information is represented by the dark grey boxes. **(a)** Wide-format data table, analogous to that in a typical VCF file. Each sample is stored in an individual column and loci are in rows. **(b)** Long-format data table, with a single column for the locus and sample identifiers. Rows thus contain every unique combination of sample and locus.

Most of GENOMALICIOUS’ functions work with around the `data.table` class objects, from R’s DATA.TABLE package (Dowle and Srinivasan 2019). There are several reasons why I believe this a very useful starting format for population genomic data. (1) Tabular data are easy to visualise and mentally interpret. (2) Tabular data in long-format is used in many R functions, allowing smoother integration of SNP data into R pipelines. (3) The `data.table` class stores large volumes of information efficiently and has excellent manipulation features. (4) Therefore, users are returned an object that they are free to manipulate to their liking. (5) The `data.table` class has some very useful manipulation features that make them ideal for dealing with large datasets; see the vignettes on CRAN for more details and explanations (https://cran.r-project.org/web/packages/data.table/vignettes/). (6) `data.table` objects are also innately dual `data.frame` objects, and can easily be converted into pure `data.frame` or `matrix` objects using the respective functions, `as.data.frame` and `as.matrix`.

## EXPORT

Once imported as a SNP `data.table`, various functions exist to convert SNP data into formats for other programs external to R. Some such functions in the current version include: (1 `bayescan_input_pools`, to export pooled allele frequencies for selection scans in BAYESCAN (Foll and Gaggiotti 2008); (2) `dadi_inputs_pools`, to construct inferred site frequency spectra from pooled allele frequencies for demographic analyses with DADI (Gutenkunst et al. 2009); and (3) `myrbayes_input`, to export an input file ready for phylogenetic analyses in MRBAYES (Huelsenbeck and Ronquist 2001; Ronquist et al. 2012).

## MANIPULATE

Several tools are provided in GENOMALICIOUS to manipulate data. There include functions available to filter SNP data, for example: `filter_depth`, `filter_maf`, and `filter_unlink`. Another functionality is the conversion of genotype or allele frequency data between long-format data tables and wide-format matrices (or vice versa): `DT2Mat_genos` and `DT2Mat_freqs`. Users can also convert genotypes into allele frequencies using `genos2freqs`, or convert between genotype scoring methods using `genoscore_converter`.

There are functions to manipulate data into more complex data structures used for programs within R. Some such functions in the current version include: (1) `adegenet_DT2genX`, to convert a `data.table` class object of genotypes into a `genind` or `genlight` class object used by ADEGENET (Jombart 2008); (2) `outflank_input_freqs`, to convert a `data.table` class object of pooled allele frequencies into a format for OUTFLANK (Whitlock and Lotterhos 2015); (3) `polyrad_DT2RADdata`, to convert a `data.table` class object of genotype read counts into a `RADdata` object used by POLYRAD (Clark et al. 2019); (4) `poolfstat_DT2pooldata`, to convert a `data.table` class object of genotypes into a `pooldata` object used by POOLFSTAT (Hivert et al. 2018).

## ANALYSE

As GENOMALICIOUS has evolved, it has included an increasing number of functions for analysis of SNPs, sequences, and genome annotations. Of interest to most population genomic investigations are the functions for examining population genetic structure. The current version provides the `fstat_calc` function to calculate *F*-statistics following (Weir and Cockerham 1984) and takes a genotype or allele frequency `data.table` class object as input. For genotypes, you can calculate *F*_ST_, *F*_IS_, and *F*_IT_ and test their significance using permutation tests. For allele frequencies, you can calculate *F*_ST_.

I also provide a series of functions to facilitate analyses of population structure using multivariate analyses. Principal component analysis (PCA) of genotype data is performed using the `pca_genos` function, which is a wrapper for R’s `prcomp` function, taking a genotype `data.table` class object as input. Principal coordinate analysis (PCoA) of allele frequency data is performed using the `pcoa_freqs` function, and is a wrapper for APE’s (Paradis et al. 2004) `pcoa` function, taking an allele frequency `data.table` class object as input.

PCA analyses of genotype data can be taken further using discriminant analysis of principal components (DAPC) (Jombart et al. 2010). Functions in GENOMALICIOUS can assist investigators in conducting these DAPC in concordance with recently described best practise guidelines (Thia 2023). Using the `dapc_infer` function, an investigator can implement a series of *K*-means clustering analyses to understand how different number of PC axes affects inference of the optimal *K* effective populations in a genotype dataset. Once an investigator has designated the groups/populations they want to discriminate, they can conduct a DAPC using the `dapc_fit` function. This function can flexibly perform a straight DAPC, or perform a cross-validation analysis using a leave-one-out approach or a training–testing partitioning approach. Such cross-validation has been highlighted as important in evaluating the fit and biological relevance of a DAPC (Thia 2023).

Aside from these core functions to analyse population structure are useful functions for more niche purposes. The `family_sim_genos`, `family_sim_compare`, and `family_sim_qtl` functions can be used to simulate family groups given a set of population allele frequencies, assess capacity to estimate relatedness coefficients, and simulate quantitative trait loci with a given additive genetic variance, respectively. The `align_many_genes_dna` and `align_many_genes_aa` functions provide wrappers for the MSA package (Bodenhofer et al. 2015), assisting the automation of multi-gene nucleotide and amino acid alignments, respectively. The `locus_overlap` function can be used to test whether sets of loci exhibit significant overlap relative to random expectations, for example, candidate loci identified in selection scans.

## VISUALISE

A spread of different functions is available in GENOMALICIOUS to visualise data. Missing data can be visualised as a heatmap using the `miss_plot_heatmap` function (Figure 2a), or as histograms using the `miss_plot_hist` function (Figure 2b and c). DNA sequence alignments can be visualised using the `align_plot_dna` and `align_plot_aa` functions (Figure 3). It is possible to produce visualisations of gene structures and organisation using the `gene_map_plot` (Figure 4). For population genetic structure, the `pcoa_plot`, `pca_plot`, and `dapc_plot` can provide visualisations of eigen vector loadings (scree plots) and ordinations of samples (scatter plots) (Figure 5). For DAPC, you can also visualise the outputs of *K*-means clustering (Figure 6a), group probabilities (Figure 6b), and cross-validation assignment rates (Figure 6c).

**Figure 2.**
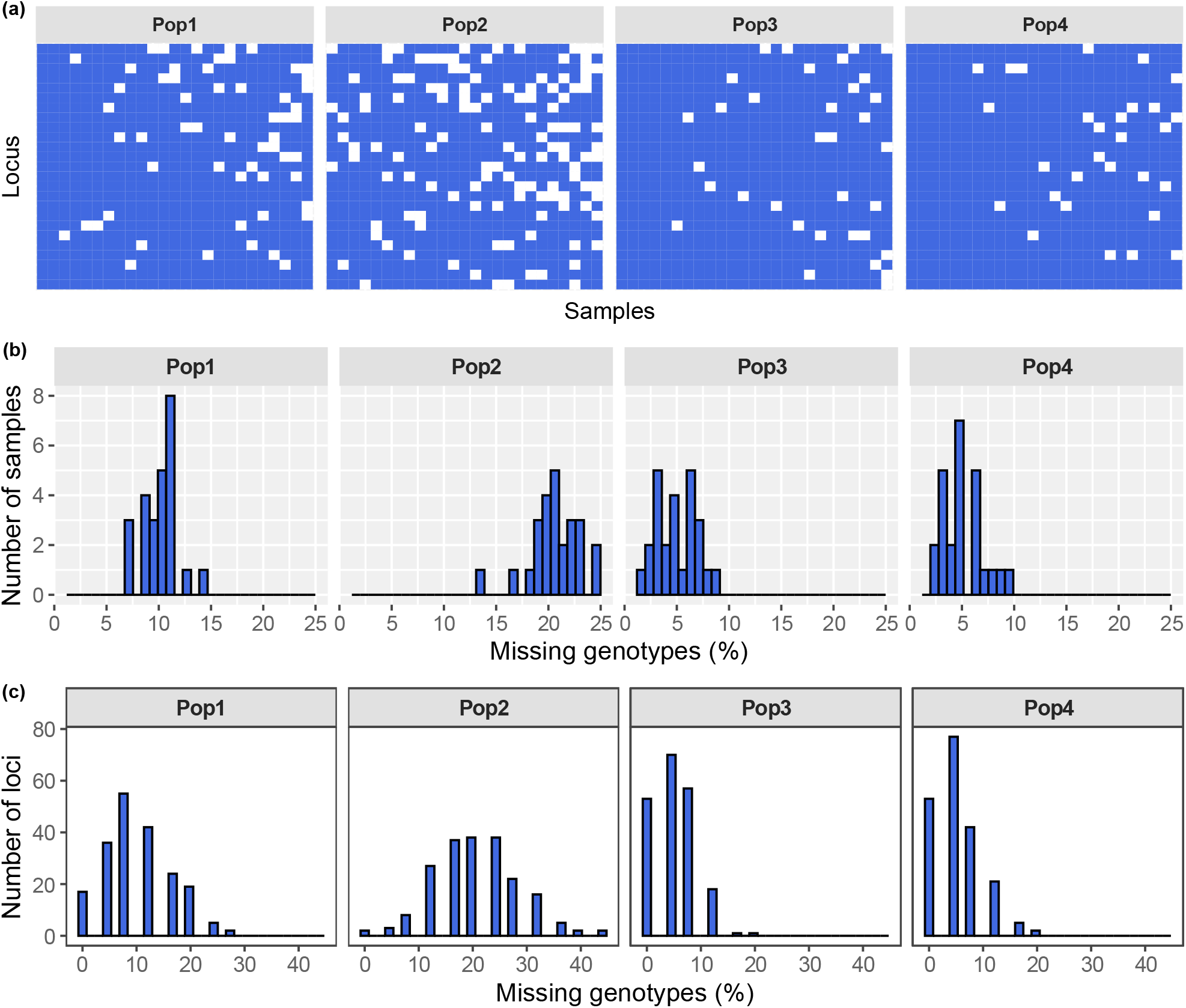
Visualisations of missing data using GENOMALICIOUS. **(a)** A heatmap of missing data. Samples (x-axis) and loci (y-axis) are plotted against each other, subset by population (panels). Colours represent the missing (white) and non-missing (blue) data for each sample and locus combination. **(b,c)** Histograms of missing data. The number of missing genotypes (x-axis) and their relative frequency (y-axis) are plotted against each other, subset by population (panels). Missing data can be plotted with respect to missingness within **(b)** samples, or **(c)** loci. The plot styles can be specified to exhibit **(b)** “ggplot” (Wickham 2016), or **(c)** “classic” R appearances.

**Figure 3.**
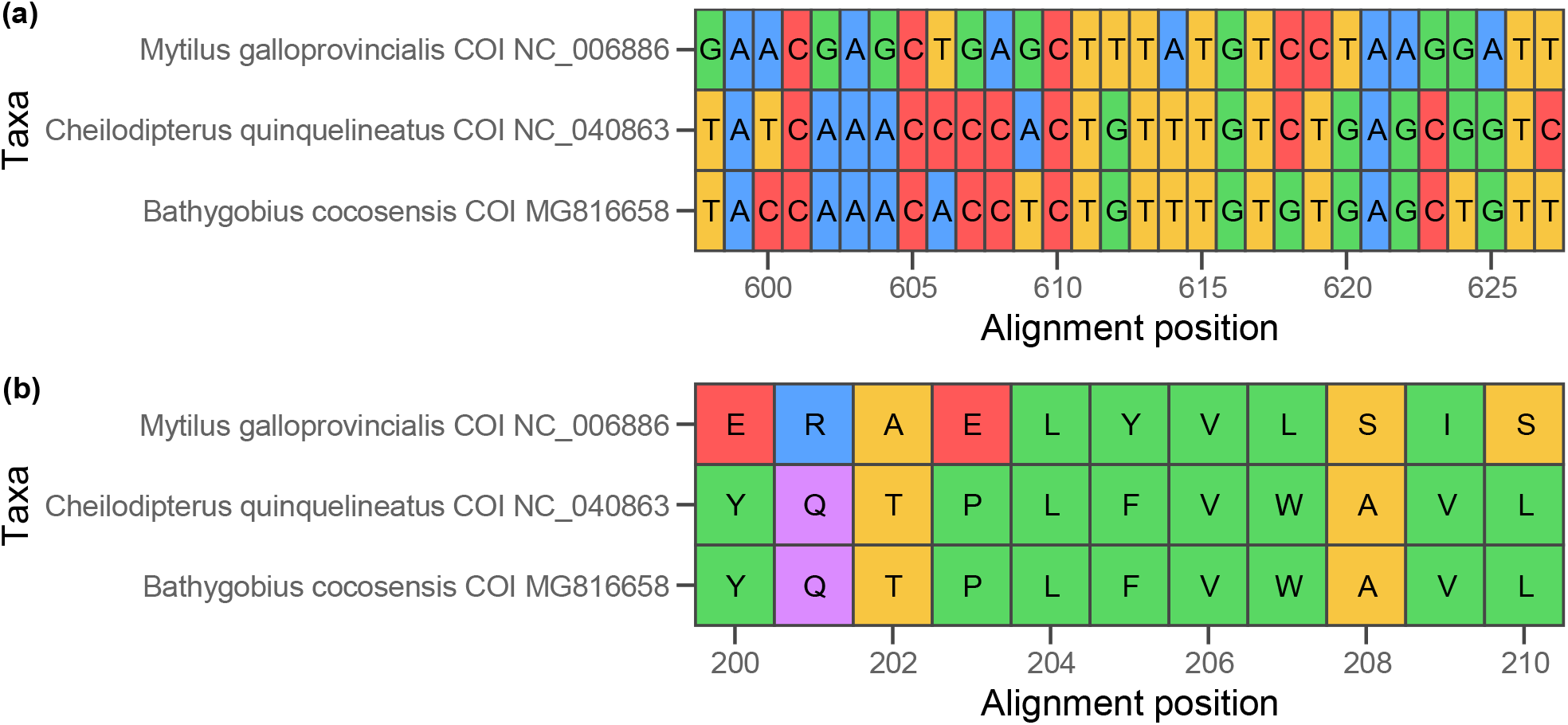
Visualisation of sequence alignments using GENOMALICIOUS. Here, a section of the COI gene is aligned for the Mediterranean mussel (*Mytilus galloprovincialis*), the five-lined cardinalfish (*Cheilodipterus quinquelineatus*) and Cocos frillgoby (*Bathygobius cocosensis*). The alignment position (x-axis) is plotted against the taxa (y-axis). Cells are coloured by their respective nucleotide or amino acid. **(a)** DNA sequence alignment from nucleotide alignment positions 598 to 627. **(b)** The corresponding amino acid sequence from residue alignment positions 200 to 210.

**Figure 4.**
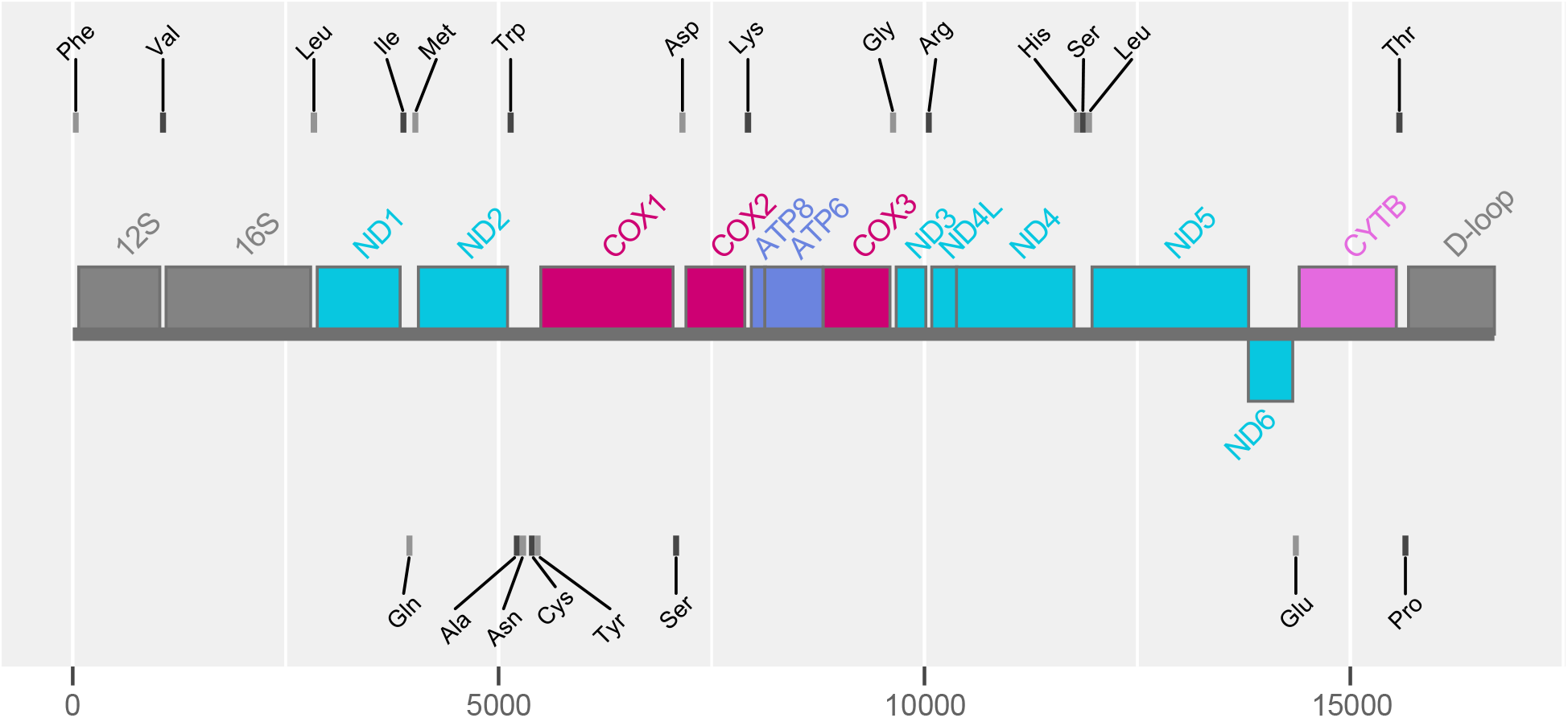
Visualisation of gene structure and organisation using GENOMALICIOUS. Here, the mitochondrial genome of Cocos frillgoby (*Bathygobius* cocosensis) is illustrated. The thin horizontal grey line represents the genome, with base positions on the x-axis. Coloured boxes sitting above and below the genome demarcate protein coding and rRNA genes. Small grey boxes floating above and below the genome demarcate tRNA coding genes. The position above versus below the genome indicates whether the gene is on the positive or negative strand, respectively. Annotations from Evans et al. (2018).

**Figure 5.**
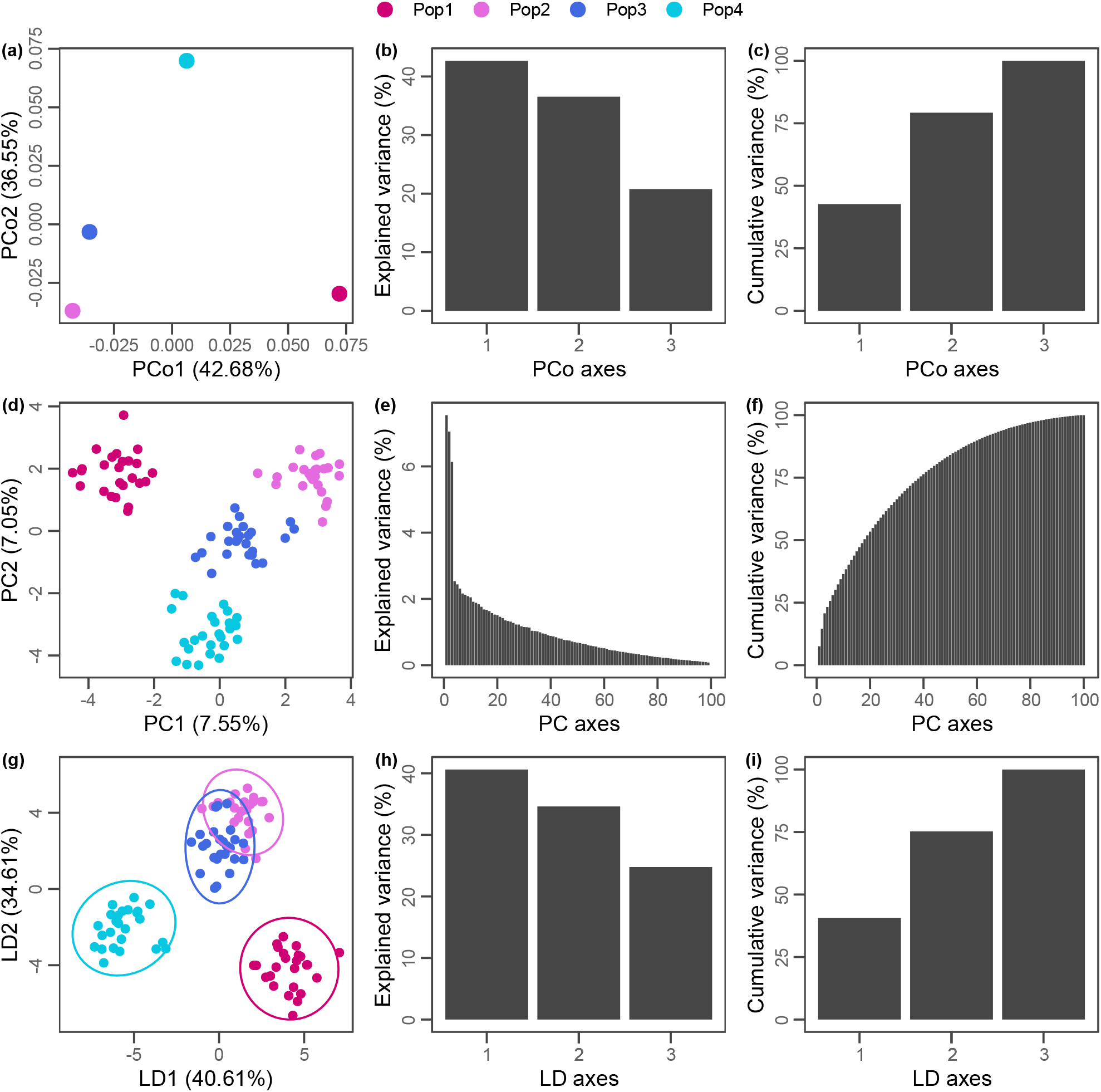
Multivariate analyses of population genetic structure using GENOMALICIOUS. **(a–c)** Results from a principal coordinate analysis (PCoA) of allele frequencies. **(d–f)** Results from a principal components analysis (PCA) of genotypes. **(g–i)** Results from a discriminant analysis of principal components (DAPC) of genotypes. **(a,d,g)** Scatterplots projecting populations or samples into multivariate space. **(b,e,h)** Screeplots of explained variance attributed to each axis. **(c,f,i)** Plots of the cumulative variance with each subsequent axis.

**Figure 6.**
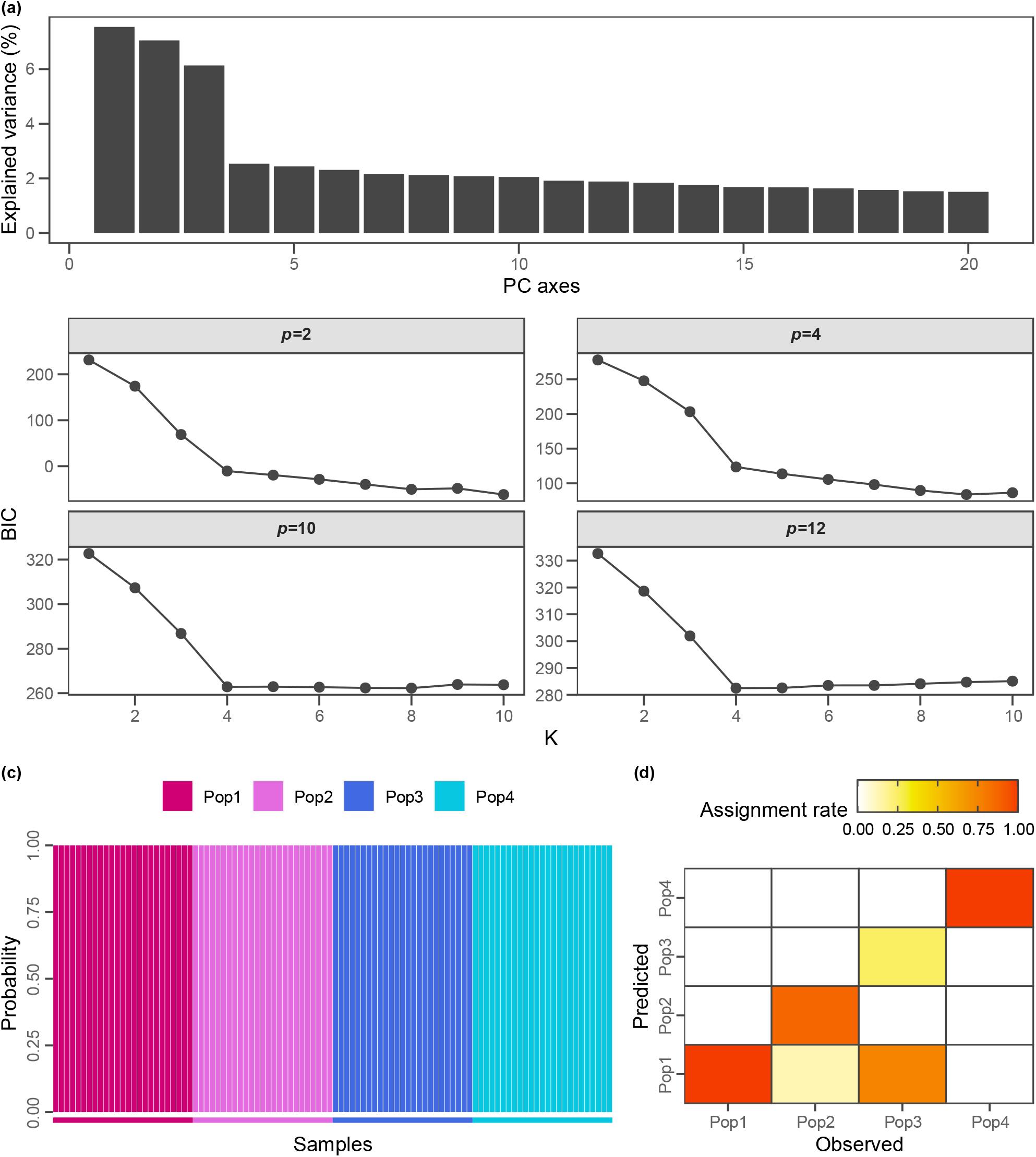
Additional functionalities in GENOMALICIOUS to facilitate best practise with discriminant analysis of principal components. **(a)** Inference of the number of *K* effective populations using *K*-means clustering. In the top row, the scree plot of explained variances for PC axes. In bottom rows, the BIC scores (y-axis) for different inferred values of *K* (x-axis) using different numbers of PC axes as predictors of among-population variation (panels) **(b)** A probability plot of posterior probability (y-axis) that a sample (x-axis) belongs to each population analysed. Designated populations are indicated by the thin coloured horizontal bars on the x-axis. Vertical bars represent individuals, with colours indicating the probability that it belongs to a given population. Here, all individuals have 100% probability of belonging to their designated population. **(c)** An assignment rate heatmap summarising the results of cross-validation analysis. The observed population (x-axis) is plotted relative to the predicted population (y-axis). Colour intensity indicates the relative assignment rate of an individual from one population to another. The diagonal indicates the correct assignment.

## SUMMARY

I developed and continue to expand GENOMALICIOUS for the purposes of providing:

1. Easy import of VCF files into R as (long-format) data tables, and import/export tools for various population genetics programs that operate external or within R.
2. Easy-to-use functions for data manipulations into commonly used population genetic data structures or more complex structures required by other functions within R.
3. Functions for basic analyses of population genomic data in R.
4. Functions for visualising population genomic data and results in R.
5. Functions that can be deployed on pooled or individually sequenced data.
6. An accessible R package for assisting the learning and teaching of population genomic analyses.

This package is an evolving entity. I look forward to feedback and any suggestions on how GENOMALICIOUS can be improved and made to better suit the appetites of population genomicists.

## ACCESSIBILITY

The most up-to-date version of GENOMALICIOUS is currently being served on J.A. Thia’s GitHub: https://github.com/j-a-thia/genomalicious. This GitHub repository also contains various tutorials to help familiarise users with GENOMALICIOUS. Code for all figures presented here is appended at the end of this manuscript.

## CONFLICT OF INTEREST DISCLOUSRE

There is no conflict of interest associated with this manuscript.

## CODE

~~~
#’ DEMO FIGURES FOR GENOMALICIOUS PAPER
#’
#’ AUTHOR: JOSHUA THIA
#’
#’ This script creates a directory called ‘Figures/’ and will write out
#’ the created figures into it. Assumes your current working directory is
#’ where you would like to run the script.
library(genomalicious)
library(ggpubr)
if(dir.exists(‘Figures’)==FALSE){ dir.create(‘Figures’) }
# >>>>>>>>>>>>>>>>>>>>>>>>>>>>>>>>>>>>>>>>>>>>>>>>>>>>>>>>>>>>>>>>>>>>>>>>>
#### 4 POPULATION DATA ####
# >>>>>>>>>>>>>>>>>>>>>>>>>>>>>>>>>>>>>>>>>>>>>>>>>>>>>>>>>>>>>>>>>>>>>>>>>
# This section takes pre-packaged population data and sets it up for
# downstream analysis.
# Population colours
pop.colours <-c(
 ‘Pop1’=‘#ce0073’, ‘Pop2’=‘#e46adf’,
 ‘Pop3’=‘royalblue’, ‘Pop4’=‘#08c7e0’
)
# Pull in genotype and allele frequency data
data(“data_Genos”)
data(“data_PoolFreqs”)
datGt <- data_Genos
datFq <- data_PoolFreqs
# Add in the number of individuals pooled to get allele frequencies
datFq[, INDS:=30]
# Convert the genotype from character to integer
datGt[, GT.INT:=genoscore_converter(GT)]
# >>>>>>>>>>>>>>>>>>>>>>>>>>>>>>>>>>>>>>>>>>>>>>>>>>>>>>>>>>>>>>>>>>>>>>>>>
#### MISSING DATA ####
# >>>>>>>>>>>>>>>>>>>>>>>>>>>>>>>>>>>>>>>>>>>>>>>>>>>>>>>>>>>>>>>>>>>>>>>>>
# Plot missing data for genotypes
datGtMiss <- do.call(
 ‘rbind’,
 # Split data table by sample, and iterate through samples, X
 split(datGt, by=‘POP’) %>%
   lapply(., function(Dpop){
     pop <- Dpop$POP[1]
     if(pop==‘Pop1’){
        pr <- 0.1
     } else if(pop==‘Pop2’){
        pr <- 0.2
     } else if(pop %in% c(‘Pop3’,’Pop4’)){
        pr <- 0.05
     }
     # Numbers and unique loci and samples
     num.loc <- Dpop$LOCUS %>%
     unique %>% length uniq.loc <- Dpop$LOCUS %>% unique
     num.samp <- Dpop$SAMPLE %>% unique %>% length
     uniq.samp <- Dpop$SAMPLE %>% unique
     # Vector of missingness
     num.miss <- rbinom(n=num.samp, size=num.loc, prob=pr)
     # Iterate through samples and add unique loci
     for(i in 1:num.samp){
       locs <- sample(uniq.loc, size=num.miss[i], replace=FALSE)
       Dpop[SAMPLE==uniq.samp[i] & LOCUS%in%locs, GT:=NA]
     }
     # Return
     return(Dpop)
     }
     )
)
# Unique loci
uniq.loci <- unique(datGtMiss$LOCUS)
# Plot
cairo_pdf(‘Figures/missing_data.pdf’, width=20/2.54, height=17/2.54)
ggarrange(
  miss_plot_heatmap(datGtMiss[LOCUS %in% uniq.loci[1:25]], sortLoci = ‘missing’,
sortSamp=‘missing’, popCol=‘POP’, plotNCol = 4),
  miss_plot_hist(datGtMiss, plotBy=‘samples’,look=‘ggplot’, popCol=‘POP’,
plotColour=‘royalblue’, plotNCol = 4),
  miss_plot_hist(datGtMiss, plotBy=‘loci’, look=‘classic’, popCol=‘POP’,
plotColour=‘royalblue’, plotNCol = 4),
  labels=c(‘(a)’,’(b)’,’(c)’),
  font.label=list(size=9, face=‘bold’),
  nrow=3, ncol=1
)
dev.off()
# >>>>>>>>>>>>>>>>>>>>>>>>>>>>>>>>>>>>>>>>>>>>>>>>>>>>>>>>>>>>>>>>>>>>>>>>>
#### COI ALIGNMENT ####
# >>>>>>>>>>>>>>>>>>>>>>>>>>>>>>>>>>>>>>>>>>>>>>>>>>>>>>>>>>>>>>>>>>>>>>>>>
# Create a link to raw external datasets in genomalicious
genomaliciousExtData <- paste0(find.package(‘genomalicious’), ‘/extdata’)
# Multi sequence alignment of CO1 DNA and amino acids for three marine
# organisms: a mussel, a cardinal fish, and a goby
co1.dna <- paste0(genomaliciousExtData, ‘/data_COI_dna.fasta’) %>%
  align_many_genes_dna(., gene.names=‘COI’) %>%
  .[[‘COI’]] %>%
  .[[‘align’]] %>%
  as.matrix()
co1.aa <- paste0(genomaliciousExtData, ‘/data_COI_aa.fasta’) %>%
  align_many_genes_aa(., gene.names=‘COI’) %>%
  .[[‘COI’]] %>%
  .[[‘align’]] %>%
  as.matrix()
# Plot nucleotide positions 598 to 627, which corresponds to residue
# positions 200 to 201.
cairo_pdf(‘Figures/co1_align.pdf’, width=17/2.54, height=8/2.54)
ggarrange(
  align_plot_dna(
     co1.dna,
     border_colour=‘grey20’,
     pos_vec=598:627,
     show_bases=TRUE
  ),
  align_plot_aa(
     co1.aa,
     border_colour=‘grey20’,
     pos_vec=200:210,
     show_residues=TRUE
  ),
  nrow=2, ncol=1, labels=c(‘(a)’,’(b)’),
  font.label = list(size=9, face=‘bold’)
)
dev.off()
# >>>>>>>>>>>>>>>>>>>>>>>>>>>>>>>>>>>>>>>>>>>>>>>>>>>>>>>>>>>>>>>>>>>>>>>>>
#### MITOGENOME MAPPING ####
# >>>>>>>>>>>>>>>>>>>>>>>>>>>>>>>>>>>>>>>>>>>>>>>>>>>>>>>>>>>>>>>>>>>>>>>>>
# Pull in the mitochondrion data for the goby, Bathygobius cocosensis.
bcocoMito <- mitoGbk2DT(‘DATA_Bcocosensis_mito.gbk’) %>%
  .[, NAME:=sub(‘tRNA-’, ‘‘, NAME)] %>%
  .[TYPE %in% c(‘tRNA’,’gene’,’rRNA’,’D-loop’)] %>%
  .[NAME==‘12S ribosomal RNA’, NAME:=‘12S’] %>%
  .[NAME==‘16S ribosomal RNA’, NAME:=‘16S’] %>%
  .[, NAME:=paste0(“‘“, NAME, “‘“)]
# Take a look at the gene names, note that they are internall quoted.
# This is a trick to make sure they are parsed properly by ggplot.
bcocoMito[TYPE%in%c(‘gene’,’rRNA’,’D-loop’)]$NAME %>% sort
# Colours for the different genes.
gene.colours <- c(
  ATP6=‘#6b84df’,
  ATP8=‘#6b84df’,
  CYTB=‘#e46adf’,
  COX1=‘#ce0073’,
  COX2=‘#ce0073’,
  COX3=‘#ce0073’,
  ND1=‘#08c7e0’,
  ND2=‘#08c7e0’,
  ND3=‘#08c7e0’,
  ND4=‘#08c7e0’,
  ND4L=‘#08c7e0’,
  ND5=‘#08c7e0’,
  ND6=‘#08c7e0’,
  ‘16S’=‘#6d6d6d’,
  ‘12S’=‘#6d6d6d’,
  ‘D-loop’=‘#6d6d6d’
)
names(gene.colours) <- paste0(“‘“, names(gene.colours), “‘“)
# Plot the mitochondrion
cairo_pdf(‘Figures/bcoco_mito.pdf’, width=17/2.54, height=8/2.54)
gene_map_plot(
  mapDT=bcocoMito,
  genome_len=16692,
  gene_colour = gene.colours,
  gene_type=c(‘gene’,’rRNA’,’D-loop’),
  extra_type=c(‘tRNA’),
  gene_txt_size = 3,
  extra_txt_size = 2.5,
  gene_size=0.2,
  gene_border=‘#595959’,
  plot_ymax=4.5,
  extra_ypos = 3
)
dev.off()
# >>>>>>>>>>>>>>>>>>>>>>>>>>>>>>>>>>>>>>>>>>>>>>>>>>>>>>>>>>>>>>>>>>>>>>>>>
#### PCOA, PCA, AND DAPC ####
# >>>>>>>>>>>>>>>>>>>>>>>>>>>>>>>>>>>>>>>>>>>>>>>>>>>>>>>>>>>>>>>>>>>>>>>>>
# Perform a principal coordinates analysis on allele frequencies
PCOA <- pcoa_freqs(dat=datFq)
# Perform a principal components analysis on genotypes
PCA <- pca_genos(dat=datGt, scaling=‘covar’, sampCol=‘SAMPLE’, genoCol=‘GT.INT’,
popCol=‘POP’)
# Perform a discriminant analysis of principal components on genotypes
DAPC <- dapc_fit(dat=datGt, pcPreds=3)
# Cross-validate the DAPC
CV <- dapc_fit(dat=datGt, method=‘train_test’, pcPreds=3)
# Plot the PCoA, PCA, and DAPC: scatter, scree and cumulative variance
cairo_pdf(‘Figures/pcoa_pca_dapc.pdf’, width=20/2.54, height=20/2.54)
ggarrange(
  pcoa_plot(PCOA, type=‘scatter’, plotLook = ‘classic’, plotColours =
pop.colours) +
    theme(axis.text.y=element_text(angle=90, hjust=0.5)),
  pcoa_plot(PCOA, type=‘scree’, plotLook = ‘classic’) +
    theme(axis.text.y=element_text(angle=90, hjust=0.5)),
  pcoa_plot(PCOA, type=‘cumvar’, plotLook = ‘classic’) +
    theme(axis.text.y=element_text(angle=90, hjust=0.5)),
  pca_plot(PCA, type=‘scatter’, plotLook = ‘classic’, plotColours = pop.colours)
+
    theme(axis.text.y=element_text(angle=90, hjust=0.5)),
  pca_plot(PCA, type=‘scree’, plotLook = ‘classic’) +
    theme(axis.text.y=element_text(angle=90, hjust=0.5)),
  pca_plot(PCA, type=‘cumvar’, plotLook = ‘classic’) +
    theme(axis.text.y=element_text(angle=90, hjust=0.5)),
  dapc_plot(DAPC, type=‘scatter’, plotLook = ‘classic’, plotColours =
pop.colours) +
    theme(axis.text.y=element_text(angle=90, hjust=0.5)),
  dapc_plot(DAPC, type=‘scree’, plotLook = ‘classic’) +
    theme(axis.text.y=element_text(angle=90, hjust=0.5)),
  dapc_plot(DAPC, type=‘cumvar’, plotLook = ‘classic’) +
    theme(axis.text.y=element_text(angle=90, hjust=0.5)),
  ncol=3, nrow=3, common.legend = TRUE,
  labels=paste0(‘(‘,letters[1:9],’)’),
  font.label = list(size=9, face=‘bold’)
)
dev.off()
# Plot a K-means test for different params in DAPC, also output a
# probability plot and cross-validation results for DAPC
cairo_pdf(‘Figures/dapc_probs_cv.pdf’, width=20/2.54, height=22/2.54)
ggarrange(
  ggarrange(
    dapc_infer(datGt, plotLook=‘classic’, pTest = c(2,4,10,12))$plot,
    labels=c(‘(a)’), font.label = list(size=9, face=‘bold’)
  ),
  ggarrange(
   dapc_plot(DAPC, type=‘probs’, plotColours = pop.colours, sampleShow = FALSE)
+
      theme(axis.text.y=element_text(angle=90, hjust=0.5)),
    dapc_plot(CV, type=‘assign’, plotColours =
c(‘white’,’#F2E900’,’#F49800’,’#F43C00’)) +
      theme(axis.text.y=element_text(angle=90, hjust=0.5)),
    widths=c(60,40), labels=c(‘(c)’,’(d)’),
    font.label = list(size=9, face=‘bold’)
  ),
  nrow=2, ncol=1, heights=c(65,35)
)
dev.off()
~~~

